# *Rag1^D600A^*, a novel catalytically inactive RAG mouse model

**DOI:** 10.1101/698332

**Authors:** Jason B Wong, Jane A Skok

## Abstract

The RAG complex (RAG1 and RAG2) can bind to recombination signal sequences of antigen receptor loci gene segments and coordinate V(D)J recombination which is the primary method of generating antigen receptor diversity. Previous biochemistry studies discovered RAG1 D600, D708 and E962 residues as essential for catalytic DNA nicking and hairpin forming activity of the RAG complex. Neutralization of each of the acidic residues does not impair DNA binding to recombination signal sequence containing DNA substrates, but cleavage of the substrates is severely compromised. These three acidic residues are thought to comprise a DDE motif that is responsible for binding to a divalent cation that is necessary for cleavage activity. Although a *Rag1*^-/-^; RAG1-D708A transgenic mouse model system has been used to study dynamics of RAG activity, transgenic expression may not precisely mimic expression from the endogenous locus. In order to improve upon this model, we created *Rag1^D600A^* mice that lack B and T cells and demonstrate a developmental block at the pro-B and DN stages, respectively. Thus, *Rag1^D600A^* mice provide a novel mouse model system for studying the poorly understood noncanonical functions of RAG1.

## Introduction

Two RAG1-RAG2 monomers form a homodimer collectively called the RAG complex. RAG can bind RSS sequences that flank V(D)J gene segments and coordinate a double stranded (ds) DNA break at the junctions between recombining B cell receptor (BCR) or T cell receptor (TCR) gene segments. Although RAG1 contains a catalytic site that executes nicking and hairpin formation, RAG2 is an essential cofactor and depletion of the latter is sufficient to cause severe immunodeficiency [1,2]. Both RAG1 and RAG2 are necessary and sufficient to complete DNA cleavage in vitro [3,4].

In order to better understand the functional components of RAG1 and RAG2, various truncations have been examined for DNA binding and cleavage activity. These studies identified a minimal core region that contains DNA binding and cleavage activity [5–7]. For RAG1, the core region contains a nonamer-binding domain, a dimerization and DNA binding domain (DDBD), a pre-RNase H domain, an RNase-H like/insertion domain and a C-terminal domain. The RAG2 core region comprises an N-terminal WD40 domain that folds into a β-propeller [8].

To further dissect the mechanisms of how RAG catalyzes DNA cleavage, highly conserved amino acids were individually analyzed in RAG1. Three important acidic residues, D600, D708 and E962 were identified. When each of these were individually mutated to a nonpolar alanine (A), DNA cleavage activity was severely compromised although RAG retained its ability to bind to the RSS of a DNA substrate [9–11]. This highlights a crucial role for these three residues in RAG cleavage activity.

DNA sequence analysis of the 600 amino acid catalytic core of RAG1 provided evidence that RAG1 evolved from DNA transposases of the Transib superfamily. Included in the sequence conservation between RAG1 and the Transib superfamily transposases is a DDE motif [12]. DDE family transposases and retroviral integrases use the DDE motif to coordinate metal cations to execute DNA nicking as a first step of catalysis [13,14]. Since D600A, D708A and E962 of RAG1 share sequence similarities to DDE motif transposases, it is tempting to speculate that RAG-mediated cleavage functions in a similar method to DDE family transposases. However, the three RAG1 amino acids are far away from each other on the linear scale.

The early crystal structures of the RAG complex revealed that it takes on a “Y” shape, where each of the two RAG1 proteins bind to each other at the base via the DDBD and RAG2 sits on top of the arms that extend up from the base [15]. Advances in structural analysis of RAG revealed that the three RAG1 acidic residues (D600, D708, E962) are located in close 3D proximity to one another [16,17]. The colocalization of the three essential acidic residues (D600, D708, E962) at the active site of RAG1 provides further support that this is a DDE motif. RAG has a strikingly similar structure to two other DDE family transposases: the well-known bacterial transposase, Tn5 and the hAT family transposase, Hermes [16,18].

The combination of early RAG1 mutational, DNA sequence and protein structural analyses strongly support the idea that D600, D708 and E962 of RAG1 comprise a DDE motif that is essential for executing RAG mediated DNA cleavage in a similar manner to DDE family transposases. RAG-mediated DNA breaks occur in two steps, nicking on one DNA strand at the junction of the RSS and gene segment, followed by hairpin formation. A H20 molecule is used for a hydrolysis reaction to produce a nick. This leaves a 3’ hydroxyl group on the first strand of DNA that can attack the phosphodiester bond on the opposite strand in a direct transesterification reaction [3]. D600, D708, and E962 are the three acidic residues that are essential for stabilizing and coordinating divalent cations, such as Mg^2+^, to direct both of these chemical reactions to initiate DNA breaks [3,8].

Neutralization of D600, D708 or E962 by mutating the aspartic acid or glutamic acid to a nonpolar alanine (A) has been shown to be sufficient to disrupt DNA cleavage activity *in vitro*, [9–11]. In order to conduct mouse experiments with catalytically inactive RAG, transgenic mice were generated that express RAG1-D708A. These early transgenic mice used BAC integrations to deliver transgenes that would have similar expression patterns to their endogenous counterpart loci. The *Rag1^-/-^;* RAG1-D708A transgenic mouse system enables the collection of B and T cells in a context where RAG can bind to DNA, but cannot cleave. This has been an important tool in many studies to investigate various properties of RAG [19–21].

Previous studies with mice harboring the HG RAG1-D708A BAC demonstrated expression patterns that coincide with lymphocyte populations undergoing V(D)J recombination in both B and T cell lineages [22]. However, when a gene is expressed outside of its endogenous location, it likely does not precisely mimic the native gene in terms of expression levels. In the case of the RAG1-D708A transgenic mice this has never been thoroughly investigated. In order to prevent the endogenous wild-type allele from expressing wild-type RAG1, it is necessary to cross the RAG1-D708A transgenic mouse onto a *Rag1^-/-^* background. Many studies use RAG1-D708A in combination with other pre-rearranged *Igh* or *Tcrb* transgenic mice, which can lead to technical complications with breeding, delays in setting up new colonies and lower efficiencies in generating mice with the desired genotype.

To circumvent these issues, we used CRISPR-Cas9 technology to make a point mutation in the endogenous *Rag1* locus to generate a RAG1-D600A catalytically inactive protein. *Rag1^D600A^* mice fail to make mature B and T cells and have developmental blocks at pro-B and DN2/3 cell stages respectively. Thus, we provide a novel mouse model in which endogenous gene expression patterns are recapitulated for a catalytically inactive form of RAG that can bind to, but cannot cleave DNA.

## Results

We used CRISPR-Cas9 to genetically engineer mutations at the endogenous *Rag1* locus to encode a RAG1-D600A protein. Briefly, mouse zygotes were injected with a mix containing: sgRNAs, Cas9 mRNAs, and a single stranded donor oligonucleotide (ssODN). Zygotes were cultured to the blastocyst stage and then transplanted into host mice. After pups were born, tail tissue was collected and founder mice were assessed by PCR. The ssODN repair template was designed to incorporate five different point mutations around the site encoding the D600 residue of RAG1. To prevent the sgRNA from cutting the DNA after repair, two silent mutations upstream of the D600 site were introduced to disrupt the PAM-sequence and sgRNA binding capability. The GAT nucleotides that encode the aspartic acid (D) at residue 600 were converted to GCC to encode an alanine (A). In order to be able to easily detect alleles with the point mutation, one additional silent mutation was designed three DNA base pairs further downstream to create a NaeI/NgoMIV restriction site. Founder mice were identified by PCR and sequenced to confirm that all the mutations from the ssODN were incorporated (**Fig 1A**). To detect *Rag1^D600A^* alleles by PCR, primers were designed to create two different sizes following restriction enzyme digest of the PCR product. Failure to digest the PCR product, partial digestion or complete digestion were respectively indicative of wild-type, *Rag1^D600A/+^*, or *Rag1^D600A/D600A^* mice (**Fig 1B**).

**Fig 1.**
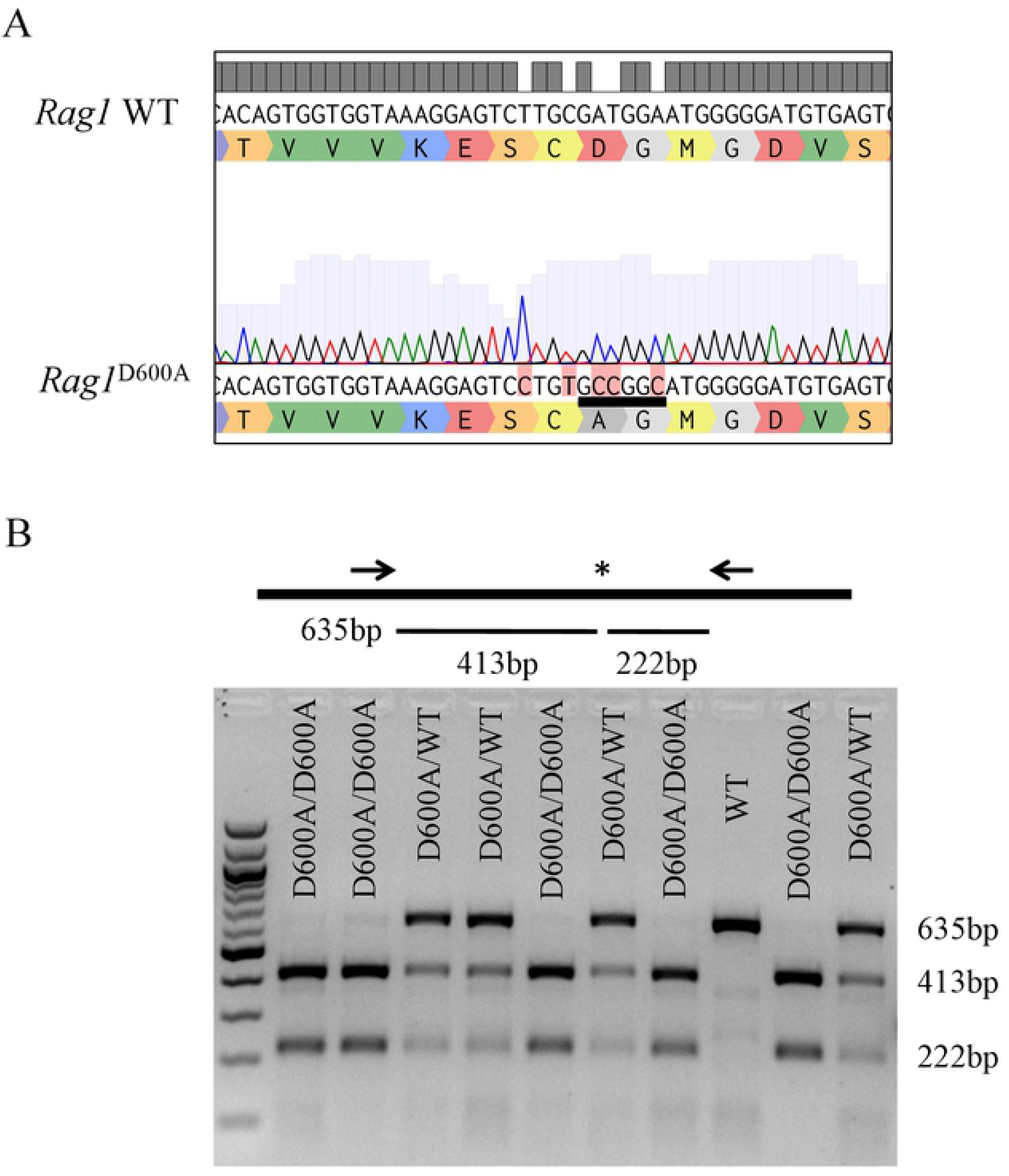
Genetic engineering for generation of *Rag1^D600A^* mice. (A) DNA sequencing of the *Rag1^D600A^* allele is shown with amino acid translation directly underneath the DNA sequence. DNA base differences highlighted in red, show mutations that disrupt secondary cutting or introduce a NgoMIV restriction site that is underlined in black. (B) genotyping PCR schematic and representative DNA gel for genotyping mice. Arrows represent PCR primers and the star shows the location of the restriction enzyme cut site. The first lane is a 100bp DNA ladder. The genotypes of mice are indicated above in subsequent lanes.

Since V(D)J recombination is essential for driving B and T cell development, it is expected that *Rag1^D600A^* mice mice will have a deficiency in B and T cells. To assess this, thymi and spleens that harbor developing T cells and mature B cells respectively were dissected from wild-type and *Rag1^D600A^* mice. The *Rag1^D600A^* thymi and spleen were decreased in size compared to wild-type littermate controls, implying a severe immune cell deficiency (**Fig 2**).

**Fig 2.**
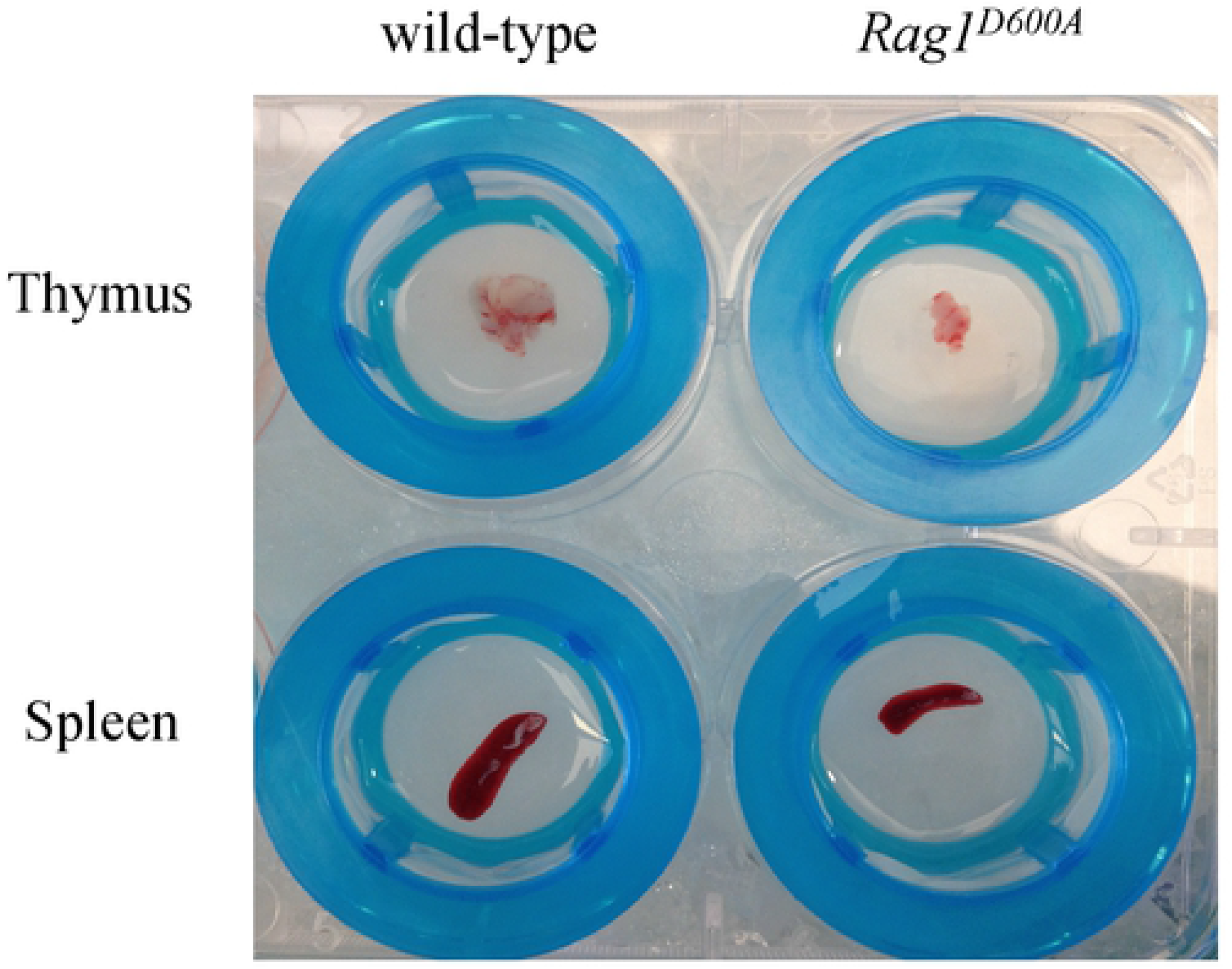
*Rag1^D600A^* mice have smaller lymphoid organs compared to wild-type. Representative images of thymi (top) and spleens (bottom) dissected from wild-type (left) or *Rag1^D600A^* (right) mice.

To better interrogate B cell deficiency in *Rag1^D600A^* mice, we performed flow cytometry analysis of developing B cells from the bone marrow. As shown in **Fig 3**, *Rag1^D600A^* mice failed to generate mature and immature B cells. Additionally, there was a block at the pro-B cell stage resulting from a failure in *Igh* recombination.

**Fig 3.**
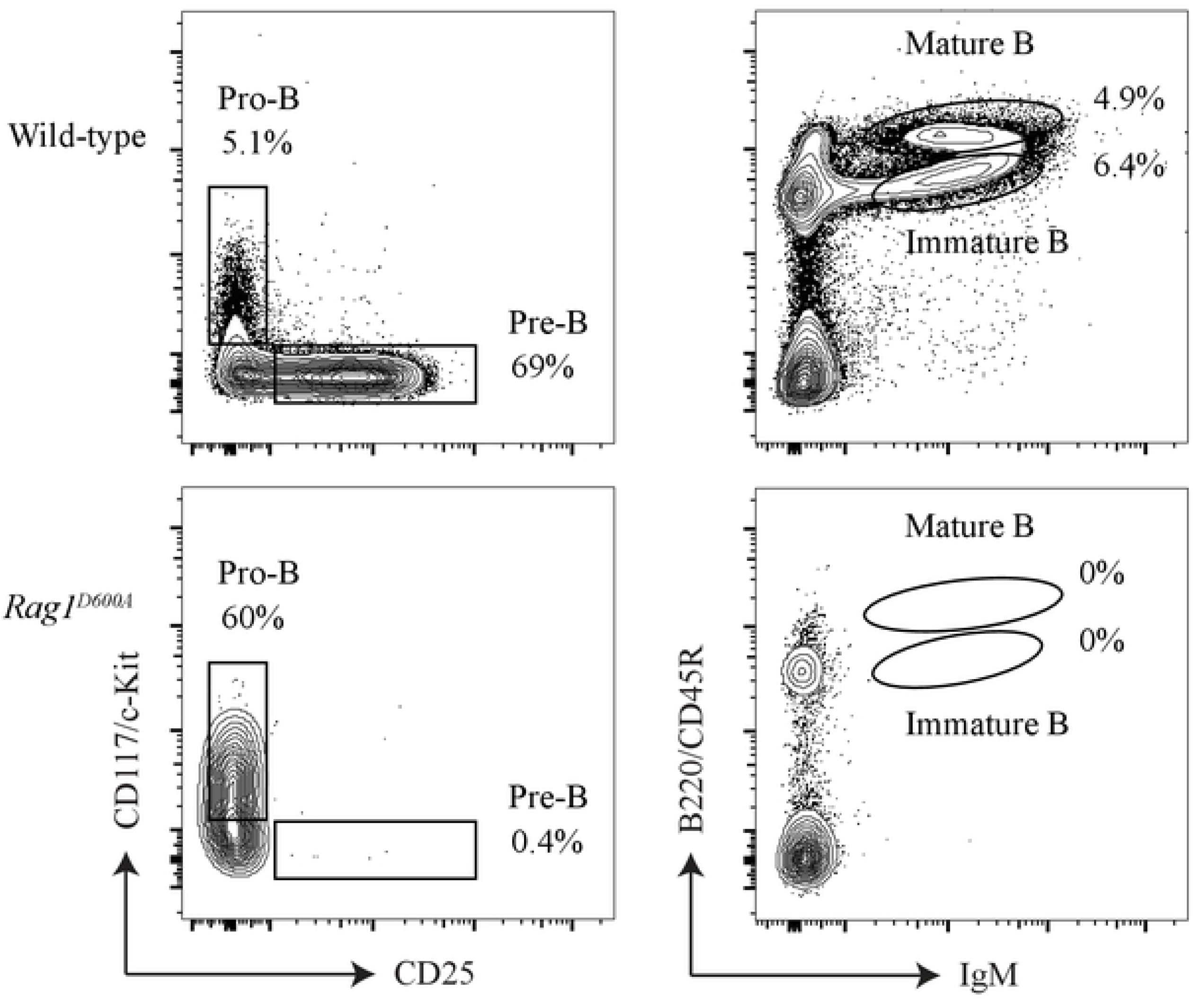
*Rag^D600A^* mice have a developmental block at the pro-B cell stage. Representative flow cytometry plots of bone marrow cells from wild-type (top) and *Rag^D600A^* mice (bottom). Left side is gated on CD19^+^, B220/CD45R^+^, IgM^-^ cells. Right side is gated on total bone marrow cells.

To determine if B cell development is also blocked at the pre-B cell stage due to a failure in *Igk* rearrangement, we crossed *Rag1^D600A^* mice with mice containing a pre-rearranged heavy-chain transgene (B1.8). *Rag1^D600A^*; B1.8 mice also failed to make mature and immature B cells, exhibiting a developmental block at the pre-B cell stage (**Fig 4**). Thus, the B1.8 allele allows B cell development to proceed to the pre-B cell stage, but development cannot proceed past the pre-B cell stage because RAG cannot recombine *Igk/Igl*.

**Fig 4.**
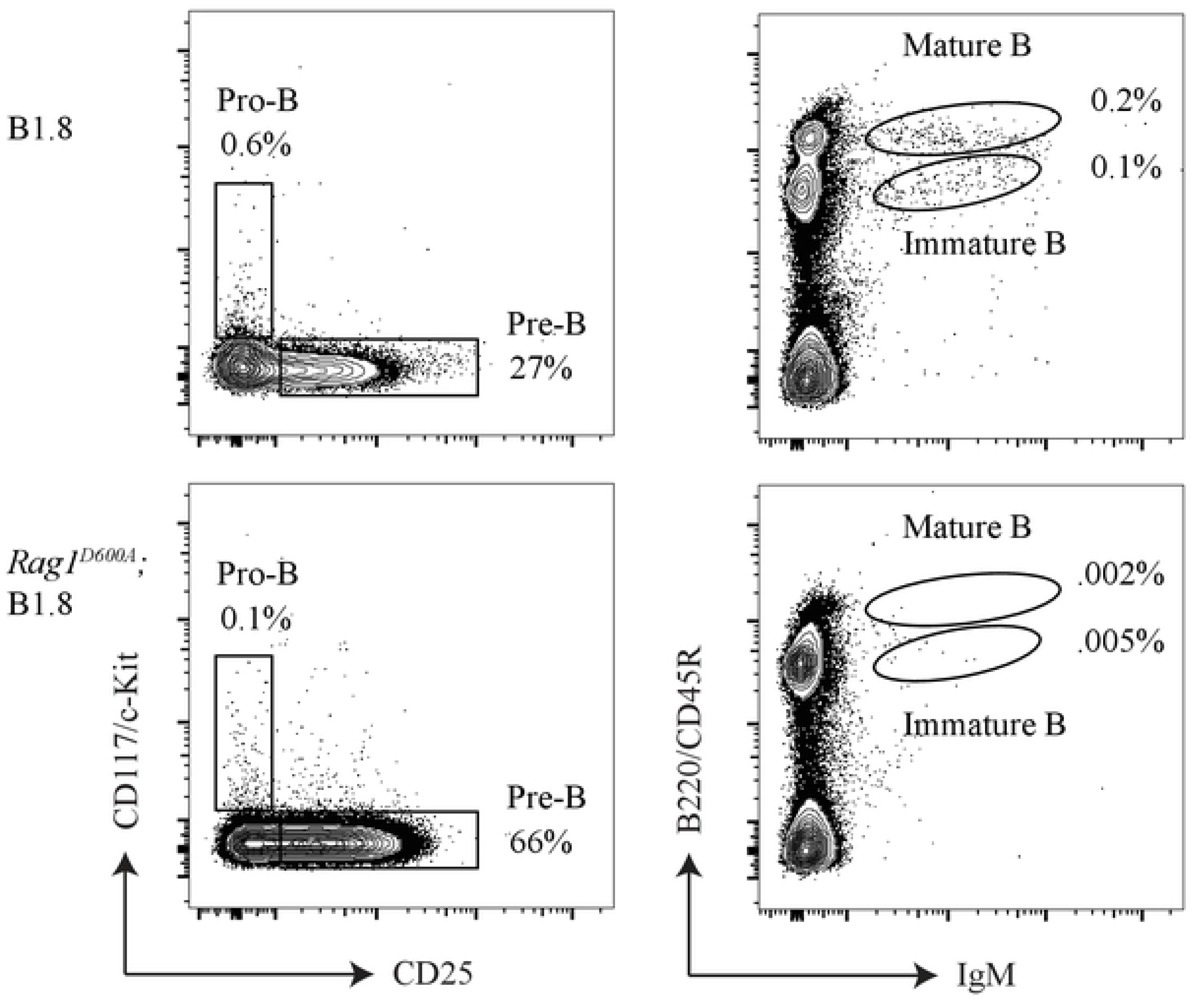
*Rag^D600A^;* B1.8 mice have a developmental block at the pre-B cell stage. Representative flow cytometry plots of bone marrow cells from B1.8 (top) and *Rag^D600A^;* B1.8 mice (bottom). Left side is gated on CD19^+^, B220/CD45R^+^, IgM^-^ bone marrow cells. Right side is gated on total bone marrow cells.

RAG is important for V(D)J recombination in both B and T cells. As expected, T cell development also appeared to be diminished based on the size of the thymus of *Rag1^D600A^* mice (**Fig 1**.). To examine this further, we analyzed T cell development from the thymus by flow cytometry. *Rag1^D600A^* T cells had a strong developmental block at the DN stage of T cell development. Additionally, these DN cells were almost exclusively (>95%) at the DN2/3 stage of development (**Fig 5**). Together our results demonstrate that *Rag1^D600A^* mice have developmental blocks in B and T cell development. Our *in vivo* findings are consistent with the previously published *in vitro* studies demonstrating that RAG1-D600A fails to catalyze DNA breaks during V(D)J recombination. Thus, *Rag1^D600A^* mice can be used as an improved mouse model for future studies.

**Fig 5.**
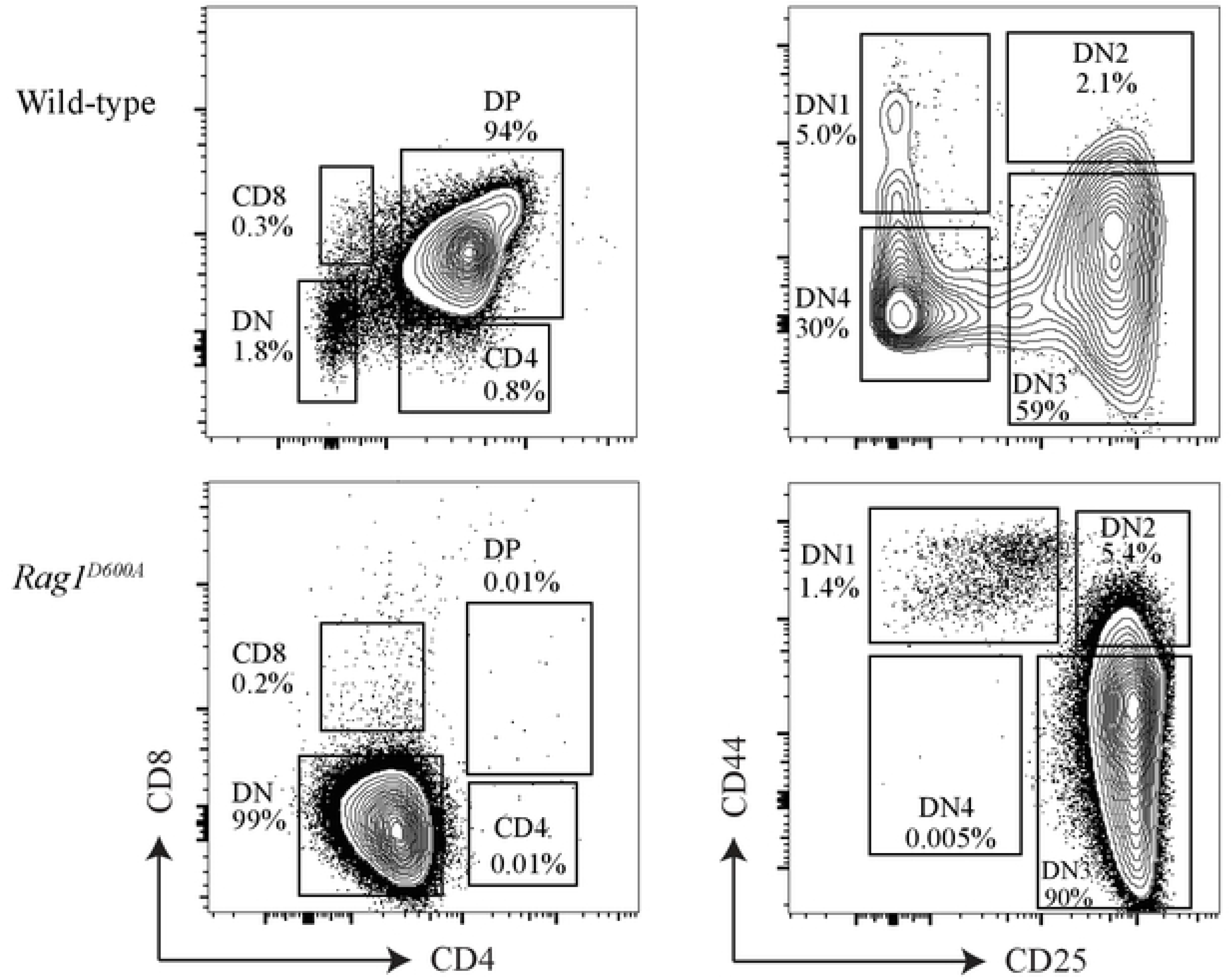
*Rag^D600A^* mice have a developmental block at the DN2/3 T cell stage. Representative flow cytometry plots of bone marrow cells from wild-type (top) and *Rag^D600A^* mice (Bottom). Left side is gated on Thy1.2^+^, TCRβ^-^ cells from the thymus. Right side is gated on the DN T cell gate.

## Discussion

Three acidic residues on RAG1 that are essential for DNA cleavage activity [9–11], D600, D708 and E962 are thought to form a DDE motif that can bind to divalent cations like Ca^2+^, Mg^2+^ or Mn^2+^ to help coordinate the chemical reactions for nicking and hairpin formation. Even though RAG1-D708A transgenic mice have been used extensively for analysis of RAG binding to DNA in the absence of cleavage, some technical issues might persist. RAG1-D708A transgenic mice express RAG1-D708A from a transgene instead of the native locus, which may not precisely mimic the expression from its endogenous location. Additionally, use of these mice require backcrossing into a *Rag1^-/-^* background to prevent *Rag1* expression from the endogenous locus. Here, we used CRISPR-cas9 to genetically engineer the endogenous *Rag1* locus to produce *Rag1^D600A^* mice.

Although we attempted to make both *Rag1^D600A^* and *Rag1^D708A^* mice, we failed to recover any founder mice that harbored the *Rag1^D708A^* allele. It is unclear why this is the case, but it is possible that the sgRNA designed for the D708 site had off target binding activity that decreased the efficiency of editing at the site of interest. Since we obtained founders for *Rag1^D600A^*, we did not continue further with the D708A design.

Early biochemistry experiments also tested cleavage activity by expressing D600C, D708C, and E962C forms of RAG1 [9–11]. Cysteine changes the divalent ion binding preference to Mn^2+^ instead of Mg^2+^ or Ca^2+^. Interestingly, all three groups who performed these experiments observed that RAG1-D708C, in the presence of Mn^2+^, was uniquely able to rescue *in vitro* cleavage activity. Since the D708C form of RAG1 was the only mutant able to rescue cleavage activity it suggested that the D708 residue is especially important, which is perhaps the reason why RAG1-D708A has been the favored form of the protein in previous studies. However, our work here demonstrates that RAG1-D600 is essential for B and T cell development. Thus, *Rag1^D600A^* is a novel mouse model that implements improvements on the *Rag1^-/-^*; RAG1-D708A transgenic system. This mouse can be used for studies on the non-canonical functions of RAG, which to-date have largely been ignored.

## Materials and Methods

### Mice

All mice were housed and cared for in accordance with IACUC guidelines and protocols approved by NYUMC (protocol #: IA15-01468). B1.8 mice were used as previously described [23].

### Generation of mutant mice using CRISPR-Cas9

Mouse zygotes were injected with the following mix: *in vitro* transcribed 50ng/uL sgRNAs, 100ng/uL Cas9 mRNA (Trilink L-6125) and 50ng/uL ssODN (IDT custom oligos). Target sequences were cloned into a Px461 plasmid using cut and paste cloning with a BbsI restriction enzyme. T7 promoter sequences were added by PCR and the PCR product was used as template to generate sgRNAs by *in vitro* transcription using the HiScribe T7 quick high yield RNA synthesis kit (NEB E2050S) and purified by an RNeasy mini kit (Qiagen 74104). Primers and oligos used are listed in S1 Table.

### Flow cytometry and antibodies

Bone marrow and fetal liver cell populations were isolated from C57Bl/6 mice via cell sorting and analyzed by flow cytometry. Antibodies for B and T cell analysis include: anti-CD45R/B220 (RA3-6B2), anti-CD19 (1D3), anti-IgM^b^ (AF6-78), anti-CD117/c-Kit (2B8), anti-CD25/IL2RA (PC61), anti-CD90.2/Thy1.2 (53-2.1), anti-TCRβ (H57-597), anti-CD8a (53-6.7), anti-CD4 (RM4-5), anti-CD44 (IM7). These antibodies were obtained from either BD Pharmigen or eBioscience. Cells were sorted using a FACSAria I (BD). Data were also collected on an LSR II (BD) and analyzed using FlowJo software.

## Acknowledgements

JAS is supported by NIH grant, R35GM122515. JBW was previously supported by the T32 CA009161 training grant (Levy) and the 2T32 AI100853-06 (Reizis) training grant. *The funders had no role in study design, data collection and analysis, decision to publish, or preparation of the manuscript*.

## Supporting Information

**S1 Table. DNA oligos used for generation of *Rag1^D600A^* mice.** Lower case letters for T7 primers are T7 promoter sequences. Lowercase letters for ssODNs represent mutations in relationship to the germline sequences.

